# HISAT: Hierarchical Indexing for Spliced Alignment of Transcripts

**DOI:** 10.1101/012591

**Authors:** Daehwan Kim, Ben Langmead, Steven L. Salzberg

## Abstract

HISAT is a new, highly efficient system for alignment of sequences from RNA sequencing experiments that achieves dramatically faster performance than previous methods. HISAT uses a new indexing scheme, hierarchical indexing, which is based on the Burrows-Wheeler transform and the Ferragina-Manzini (FM) index. Hierarchical indexing employs two types of indexes for alignment: (1) a whole-genome FM index to anchor each alignment, and (2) numerous local FM indexes for very rapid extensions of these alignments. HISAT’s hierarchical index for the human genome contains 48,000 local FM indexes, each representing a genomic region of ~64,000 bp. The algorithm includes several customized alignment strategies specifically designed for mapping RNA-seq reads across multiple exons. In tests on a variety of real and simulated data sets, we show that HISAT is the fastest system currently available, approximately 50 times faster than TopHat2 and 12 times faster than GSNAP, with equal or better accuracy than any other method. Despite its very large number of indexes, HISAT requires only 4.3 Gigabytes of memory to align reads to the human genome. HISAT supports genomes of any size, including those larger than 4 billion bases. HISAT is available as free, open-source software from http://www.ccb.jhu.edu/software/hisat.

## Background

Since its introduction in 2008, high-throughput RNA sequencing technology (RNA-seq)^1^ has become ubiquitous as a tool for the study of gene expression. It has been successfully applied to a variety of scientific questions including determining the structure of transcripts, quantifying expression changes, identifying long non-coding RNAs, and discovering fusion genes, among others^2^^-^^5^. As RNA-seq has matured, sequencing throughput and read lengths have increased dramatically: in 2008 a single RNA-seq run would comprise several million reads approximately 25-40 bp long, while today a run might contain 100-500 million reads with lengths of 100 bp or longer. These large and ever-increasing data volumes necessitate fast and scalable computational analysis systems.

RNA-seq analysis begins by aligning reads against a reference genome to determine the location from which the reads originated^6^^-^^8^. Because of the enormous data volumes involved, this alignment step has grown so time consuming that it has become a critical bottleneck; for example, widely-used alignment programs such as TopHat2^9^ and GSNAP^10^ can take several days to process a single RNA-seq experiment. It is now common to analyze dozens of samples, and some projects are collecting much larger data sets, with hundreds or even thousands of samples, which some currently available programs require weeks or months to process. The recently introduced STAR program^11^ uses suffix arrays to provide significant faster processing than most other methods, including TopHat2. However, the suffix array method has very large memory requirements (28 GB for the human genome) and as we show later, yields substantially lower alignment sensitivity than the best methods.

To create a much faster spliced aligner that uses a modest amount of RAM, we designed a novel indexing strategy, which we call *hierarchical indexing*, and implemented it in a new program, HISAT (Hierarchical Indexing for Spliced Alignment of Transcripts). The new indexing scheme is based on the Burrows-Wheeler transform^12^ and the FM index^13^, which together allow for extremely fast DNA sequence alignment with a low memory footprint. HISAT uses the Bowtie2^14^ implementation to handle many of the low-level operations required to construct and search an FM index. In contrast to most other aligners, our algorithm employs two different types of indexes, as illustrated in Figure 1: (1) a global FM index that represents the entire genome, and (2) numerous small FM indexes for regions that collectively cover the genome, where each index represents 64,000 bp. For the human genome, we create ~48,000 local FM indexes, each overlapping its neighbor by 1,024 bp, to cover the entire 3 billion bases. The overlapping boundaries make it easier to align reads that would otherwise span the regions covered by two indexes.

**Figure 1.**
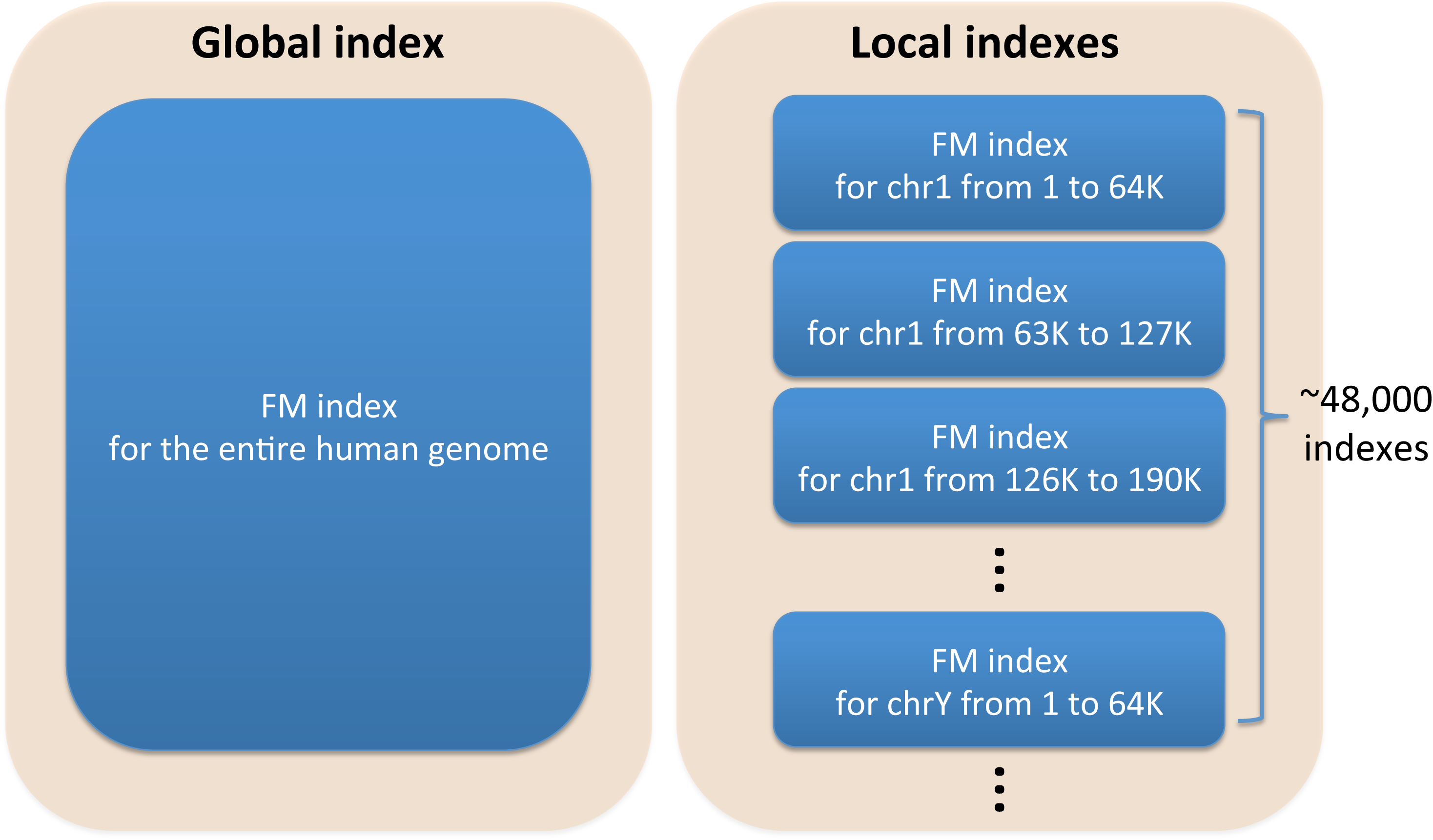
Hierarchical Indexing. Hierarchical indexing consists of two types of indexes: (1) one global index that represents the entire human genome and (2) ~48,000 local indexes that collectively cover the genome. Both types of indexes are FM-indexes, which enable extremely fast searches with a low memory footprint.

Although the program uses a very large number of local indexes, we store them in a small set of files and implement other optimizations to minimize the memory requirements, allowing us to index the human genome in approximately 4 GB of space. As we show below, hierarchical indexing enables very fast and sensitive alignment of RNA-seq reads, including multi-exon spanning reads, which other methods have difficulty aligning.

In contrast to alignment of DNA-seq reads, the alignment of RNA-seq reads comes with two additional challenges. One is that RNA-seq reads may span large gaps corresponding to introns, which in mammalian genomes can be over a megabase in length. Exons are relatively short, and thus when using 100-bp reads, a significant proportion of reads (~34.4% in our simulated data set; see Results) will span two exons. For the purpose of alignment, we divide these exon-spanning reads into three categories (Figure 2a): long-anchored reads, which have at least 15 bp in each of the two exons; intermediate-anchored reads, which have 8-15 bp in one exon; and short-anchored reads, with just 1-7 bp aligned to one of the exons.

**Figure 2.**
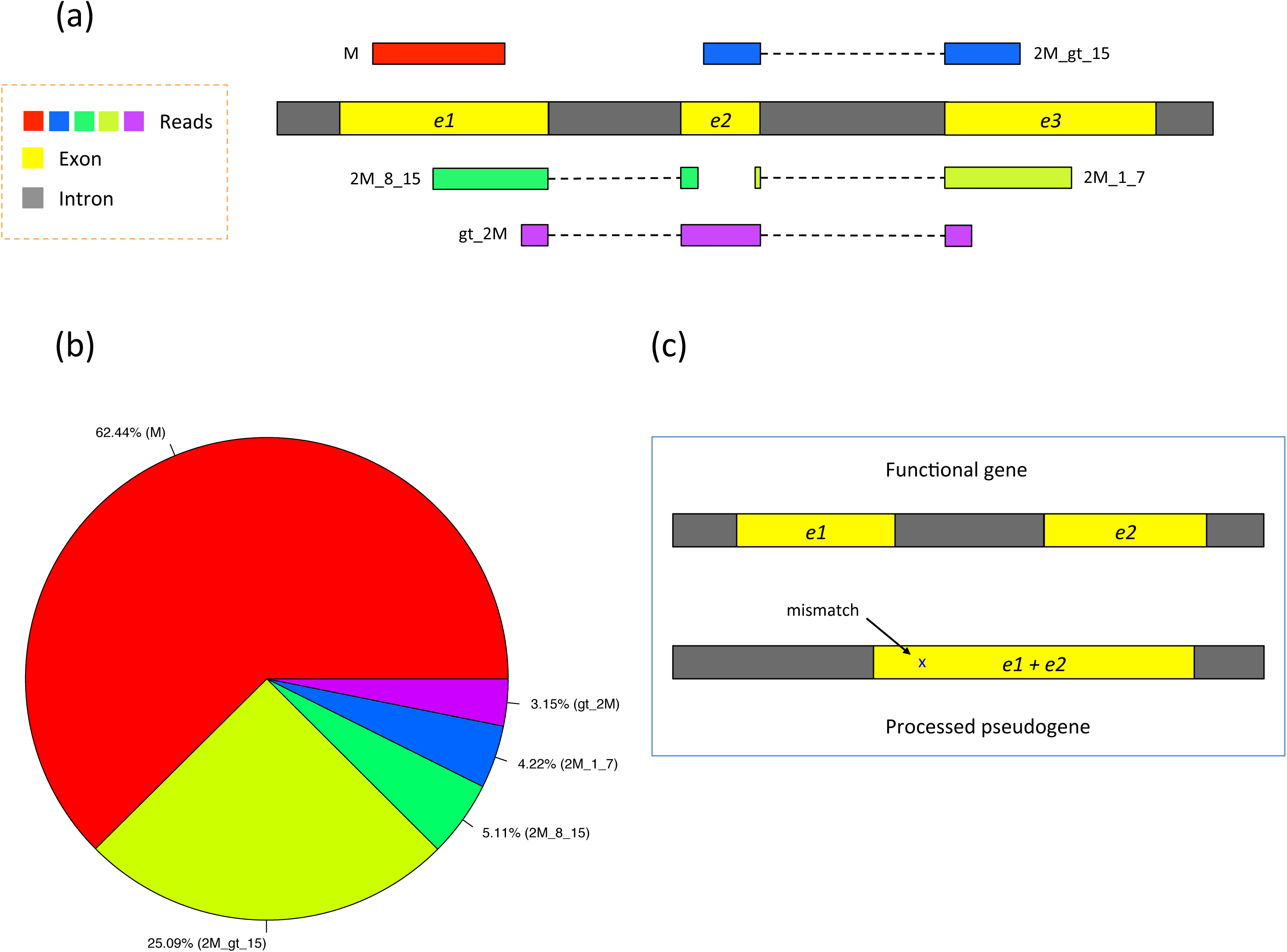
RNA-seq read types and their relative proportions from 20 million simulated reads of 100-bp long. In (a) of this figure five types of RNA-seq reads are shown: (1) M (exonic read): reads from only one exon, that is, they do not span two or more exons. M stands for exon. (2) 2M_gt_15 (junction reads with long anchors): reads spanning two exons with anchors greater than 15 bp on both exons. 2M stands for spanning two exons. (3) 2M_8_15 (junction reads with intermediate anchors): reads spanning two exons with the smaller anchor between 8 and 15 bp long. (4) 2M_1_7 (junction reads with short anchors): reads spanning two exons with the smaller anchor between 1 and 7 bp long. (5) gt_2M (junction reads spanning greater than two exons). In (b), the pie chart shows the relative proportions of different types of reads from 20 million simulated reads (100-bp long). In (c), a functional gene is depicted with its non-functional copy (processed pseudogene). Note that the pseudogene is almost identical to its parent gene, with the intron absent and one base difference in this specific example.

Figure 2b shows the proportion of reads expected to fall in each of these categories for 100-bp reads. Using simulated human RNA-seq data with realistic parameters, ~25% of the reads span two exons with long anchors (> 15 bp) in both exons, denoted in the figure by `2M_gt_15`. These reads are relatively easy to align because both anchors are long enough to be mapped to a unique location on average in the human genome. 5.1% of the reads span two exons with an intermediate-length anchor (8-15 bp) on one exon. Because most alignment programs rely on a global index, they have great difficulty mapping these reads because 8-15 bp segments will align to far too many locations (e.g., an 8-bp sequence is expected to occur ~48,000 times in the human genome). This is where the use of a local index provides a substantial advantage. In HISAT, each local index covers 64,000 bp, which ensures that over 90% of the annotated introns in the human genome are completely contained in one local index. This in turn means that HISAT can usually align small anchors using only a single local index, rather than searching across the whole genome. On average, an 8-bp sequence will occur just once in a local index of this size.

In our sample data, 4.2% of the reads span two exons with a very short anchor (1-7 bp) in one exon. Because these anchors are so short, the best approach is, where possible, to align these reads by making use of splice site information found by aligning other reads in the same data, or by using known splice sites. Note that ~3.2% of reads span more than two exons, denoted in Figure 2 by `gt_2M` (greater than two exons). In many mapping algorithms, the alignment of short- and intermediate-anchored reads and reads spanning more than two exons (12.5% of the total reads) takes up to 30-60% of the total runtime, and many of those reads are ultimately aligned incorrectly or left unaligned.

A second challenge is that processed pseudogenes, which are non-functional copies of genes and lack introns, often misdirect the mapping of reads, as illustrated in Figure 2c. Reads are often incorrectly mapped to pseudogenes instead of the genes from which they originated. This is a significant problem for the human genome, both because it contains over 14,000 pseudogenes and because genes that have pseudogene copies tend to be abundantly expressed^9^.

HISAT solves these aforementioned problems using hierarchical indexing and several alignment strategies specifically designed for handling different read types, as described in the Results and the Methods sections. HISAT is open-source software freely available for downloading at http://www.ccb.jhu.edu/software/hisat.

While HISAT is the first system to employ a hierarchical indexing strategy for spliced alignment, the strategy itself could be adopted by other methods, if their data structures can be suitably re-designed. All the programs that were included in our study – GSNAP, STAR, OLego^15^, and TopHat2 – could in principal use hierarchical indexing and thereby improve their alignment speed and quality.

## Results and discussion

We compared the accuracy and speed of HISAT to several of the leading spliced alignment programs, including STAR, GSNAP, OLego, and TopHat2, using both simulated and real reads. We tested three versions of HISAT (HISATx1, HISATx2, and HISAT), which we ran with different parameters (see Supplemental material for details). HISATx1 uses a one-pass approach that aligns each pair of reads independently of others. The second version, HISATx2, is a two-pass version of HISAT, to mimic the two-step approach used in TopHat2. In this version, the first run reports a list of splice sites supported by reads with long anchors. The second run makes use of that splice site information to align reads with short anchors (see Methods). As expected, HISATx2 takes twice as long to run, but it discovers more alignments. The STAR program also has a two-pass mode, denoted here as STARx2, which we included in our evaluation. We found that STARx2 was more than twice as slow as STAR’s default one-pass mode. This is because, prior to its second pass, STAR must build a new index for the splice junctions found in the first pass. Index building is relatively slow compared to alignment, and adds significantly to the overall time.

The third variant of HISAT (simply called HISAT, because this is the default version of the program) combines the first two ideas to gain sensitivity without the large performance cost incurred by running the program twice. In this algorithm, we allow HISAT to make use of splice sites found during the alignment of earlier reads when aligning later reads in the same run. As we show in our results below, this hybrid approach finds almost all the alignments found by HISATx2, with runtime nearly as fast as HISATx1. To the best of our knowledge, this hybrid approach is the first such singlepass method that bypasses the time-consuming step of remapping reads but matches the sensitivity of two-pass methods. HISAT also includes an option to use known splice sites from gene annotations.

For our simulated data sets, we generated 20 million reads, each 100 bp long, from 17,582 randomly chosen transcripts from known protein-coding genes, based on the GRCh37 assembly of the human genome. Each transcript was assigned expression values according to a model provided by the Flux Simulator^16^ (see Supplementary material for more details). Because we know the true alignments for the simulated reads, we can calculate alignment sensitivity as well as the sensitivity and precision of splice site detection for each program. In the results here, we discuss the performance of HISAT and other programs on error-free reads. We also ran all programs on a simulated data set that included sequencing errors. These results, which are consistent with the results on error-free data, are shown in Figure S3 and Table S3.

Figure 3 shows the alignment speed of the programs for all reads. Speed is shown as the number of reads processed per second (rps). HISATx1 and HISAT were fastest, at 141,259 and 132,781 rps respectively, and STAR was third fastest at 103,470 rps. As expected, HISATx2 (68,640 rps) and STARx2 (51,598 rps) took approximately twice as long as HISATx1 and STAR, respectively. Note that the speed reported for STARx2 did not include the index-building time, which took approximately one hour for this data set. GSNAP was substantially slower at 24,891 rps, and the slowest programs were TopHat2 (2,092 rps) and OLego (849 rps).

**Figure 3.**
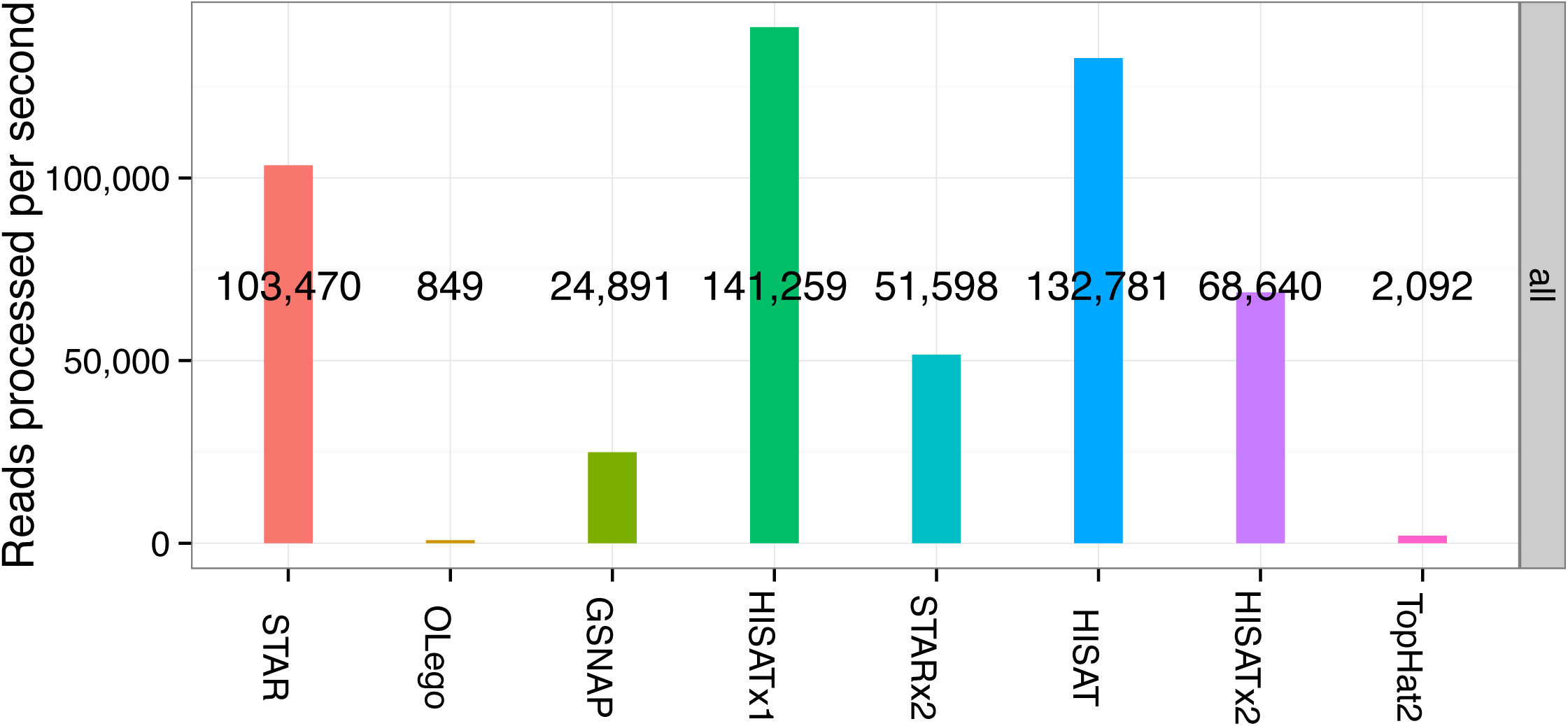
Alignment speed of spliced alignment software for 20 million simulated 100-bp reads. This figure shows the alignment speed for all the reads (M, 2M_gt_15, 2M_8_15, 2M_1_7, and gt_2M) in terms of the number of reads processed per second. Please see Figure S1 for the alignment speed for each type of read separately.

The current version of GSNAP uses a suffix array in addition to its use of a 15-mer hash table, which makes it several times faster than earlier versions that used only the hash table. OLego aligns reads using a global index based on an FM index, similar to HISAT’s algorithm. However it runs very slowly, presumably because of implementation details. Overall for this simulated data set, HISATx1 is 37% faster than STAR and six times faster than GSNAP, 68 times faster than TopHat2, and 166 times faster than OLego. As we show below, HISAT is more accurate and is only 6% slower than HISATx1.

Figure 4 shows alignment sensitivity for all programs on the 20 million simulated reads. Alignment sensitivity measures the percentage of reads that are aligned correctly, where the beginning, end, and all GT/AG splice sites within the alignment must match precisely. For non-GT/AG splice sites, an alignment was counted as correct if the intron boundaries matched within a 5-bp window. (Note that non-consensus splice sites occur in just 0.6% of all reads. Table S1 provides separate accuracies on this subset of splice sites when they are required to match precisely.) Among the one-pass algorithms (HISATx1, STAR, GSNAP, and OLego), HISATx1 and GSNAP provide the highest alignment sensitivity at 94.1%. OLego and STAR yielded lower sensitivity, at 92.2% and 91.5%, respectively.

**Figure 4.**
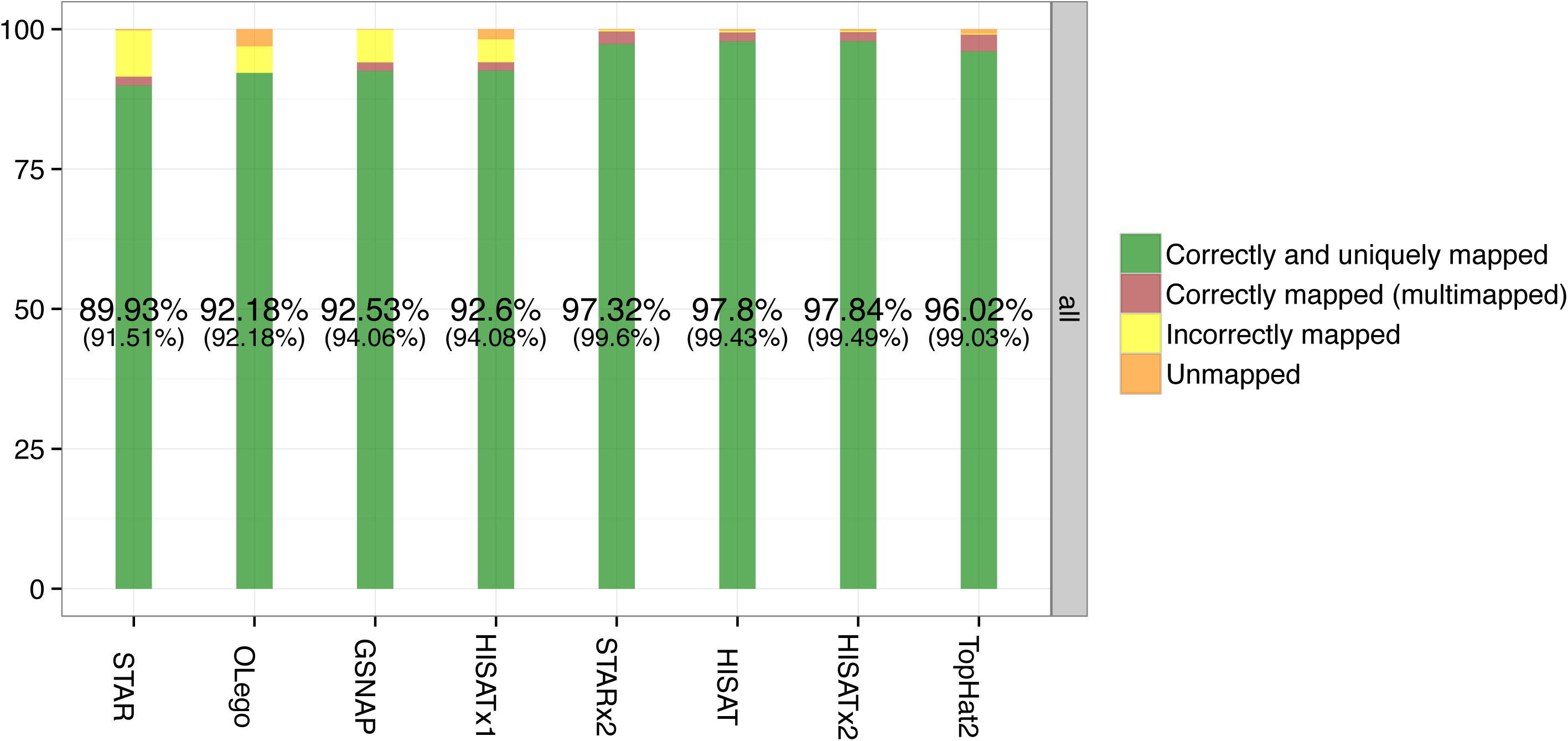
Alignment results of spliced alignment software for 20 million simulated 100-bp reads. This figure shows the alignment results for all the reads (all, M, 2M_gt_15, 2M_8_15, 2M_1_7, gt_2M). Reads are categorized as one of (1) correctly and uniquely mapped, (2) correctly mapped (multi-mapped), (3) incorrectly mapped, and (4) unmapped. Case (2) covers instances where an aligner mapped a read to multiple locations and one of the locations was correct. These four categories encompass all of the reads. The numbers in the figure represent the percentages of case (1). The numbers inside the parentheses represent the percentages of cases (1) and (2) combined.

Compared to the one-pass programs, two-pass approaches (HISATx2, STARx2, TopHat2) and the hybrid approach of HISAT obtain higher overall accuracies. These methods are much better at aligning reads with short and intermediate-length anchors. These four methods obtained sensitivity from 99.0 to 99.6%, which was 5% better than HISATx1 or GSNAP.

Figures 5 and S1, which show alignment sensitivity for reads with shorter anchors, reveal much more dramatic differences among the aligners. The two-pass algorithms (HISATx2, STARx2, TopHat2) and HISAT generate much better alignment sensitivity for short-anchored reads (1-7 bp anchors) and for reads spanning more than two exons. For reads with intermediate length anchors (Figure S1), HISATx2, STARx2, HISAT, and TopHat2 each correctly aligned >97.7% of the reads, while the one-pass methods ranged from 58.7% to 94.6%. For the reads with the shortest anchors, HISATx2, STARx2, HISAT, and TopHat2 all provided sensitivity higher than 97.5%, while the other aligners correctly aligned fewer than 10% of these reads.

**Figure 5.**
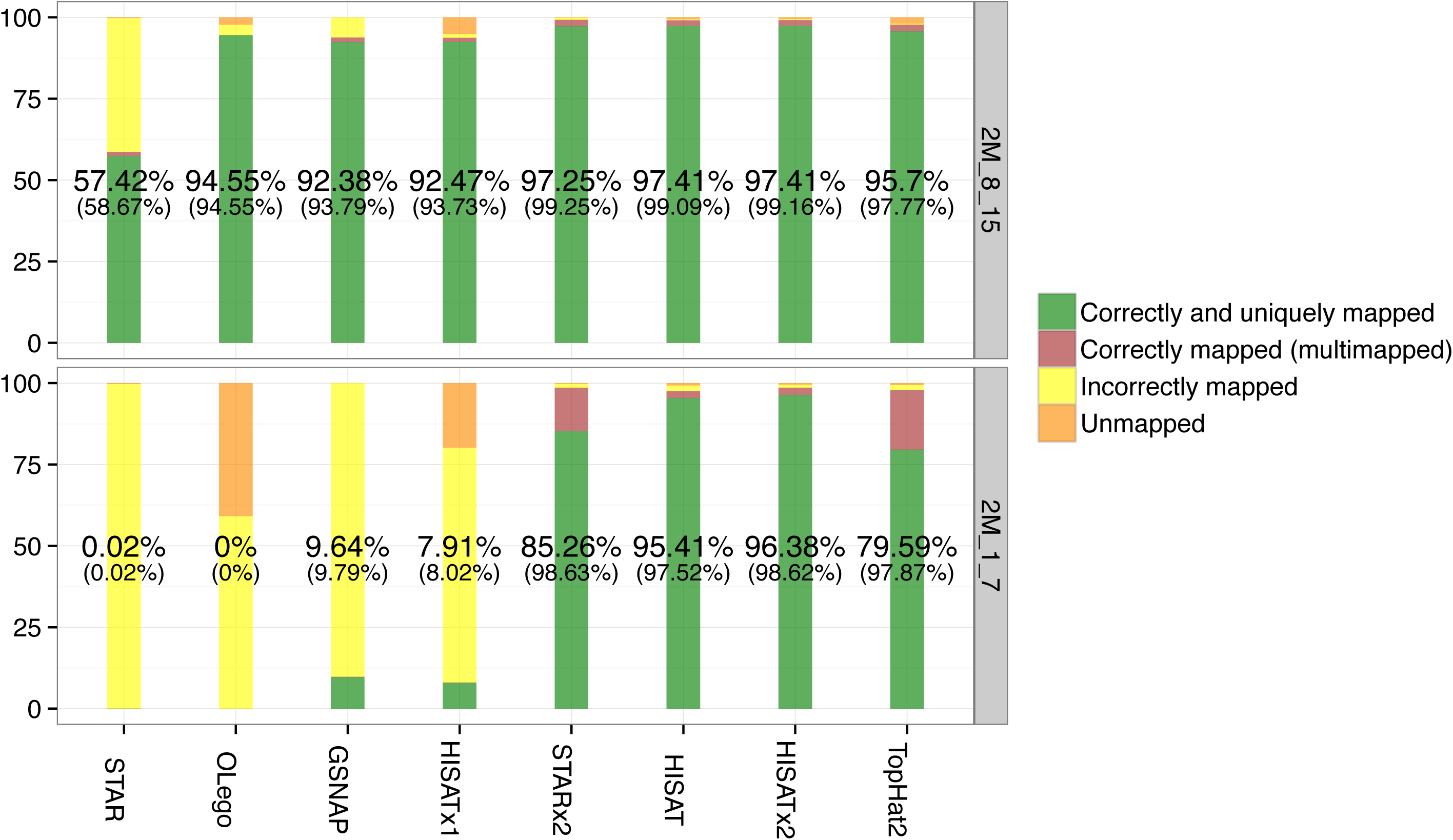
Alignment results of spliced alignment software for reads with small anchors (2M_8_15 and 2M_1_7) from 20 million simulated reads. This figure shows the alignment sensitivity for reads with small anchors (2M_8_15 and 2M_1_7). Reads are categorized as one of (1) correctly and uniquely mapped, (2) correctly mapped (multi-mapped), (3) incorrectly mapped, and (4) unmapped. Case (2) covers instances where an aligner mapped a read to multiple locations and one of the locations was correct. These four categories encompass all of the reads. The numbers in the figure represent the percentages of case (1). The numbers inside the parentheses represent the percentages of cases (1) and (2) combined. There are 1,021,935 and 843,959 reads in 2M_8_15 ad 2M_1_7, respectively.

This analysis reveals one of the key weaknesses of STAR: it aligned only 58.7% of the intermediate anchored reads, and it aligned almost none (< 0.1%) of the short-anchored reads (those with only 1-7 bp aligned to one of the exons). OLego had better sensitivity (94.6%) for intermediate-anchored reads, but it failed to align any reads with 1-7 bp anchors, and it is more than 1200 times slower than HISATx1 (see Figure S1). This illustrates the great difficulty in aligning reads with small or very small anchors using only a global index: these short anchors can be mapped to very large numbers of locations in the human genome.

We separately calculated accuracy at detecting splice sites, shown in Table 1. The simulated reads contained a total of 87,637 pairs of splice sites (acceptor sites and donor sites). We asked how many of these sites were correctly detected by each program, and gave a program credit if at least one alignment supported a given splice site. We defined precision, or positive predictive value, as the percentage of predicted sites that matched a true splice site. By these measures, HISAT obtained the highest sensitivity (97.6%) and the second highest precision (96.6%) among all the aligners. OLego yielded slightly higher precision (97%), but at the expense of lower sensitivity (94.1%).

**Table 1.** Sensitivity and precision of leading spliced aligners for 87,637 true splice sites contained in 20 million simulated reads from the human genome. Sensitivity is the percentage true splice sites found out of the total that were present. Precision (or positive predictive value) is the percentage of reported splice sites that are correct.

| Program | No. of splice sites reported | No. of true splice sites reported | Sensitivity (%) | Precision (%) |
| --- | --- | --- | --- | --- |
| HISATx1 | 89,793 | 85,466 | 97.52 | 95.18 |
| HISATx2 | 88,512 | 85,527 | 97.59 | 96.63 |
| HISAT | 88,488 | 85,503 | 97.56 | 96.63 |
| STAR | 89,717 | 84,722 | 96.67 | 94.43 |
| STARx2 | 89,116 | 84,769 | 96.73 | 95.12 |
| GSNAP | 89,486 | 85,239 | 97.26 | 95.25 |
| OLego | 84,969 | 82,428 | 94.06 | 97.01 |
| TopHat2 | 93,752 | 79,516 | 90.73 | 84.82 |

As a test on real data, we compared the aligners using 108,749,331 101-bp RNA-seq reads collected from fetal lung fibroblasts (GEO accession GSM981249; see Supplementary material for more details). Because we do not know the true alignments for these reads, we cannot tell which programs placed them at the correct locations on the genome. However, we can evaluate alignment quality in two other ways. We considered the following two criteria: (1) the cumulative number of alignments detected, up to a given edit distance, for edit distances from 0 to 3; and (2) the number of spliced alignments found that correspond to known human splice sites, based on the Ensembl GRCh37 gene annotation.

Figure 6 shows the cumulative number of alignments found by each program on this data set, divided according to edit distances from 0 to 3. Edit distance is defined here simply as the number of differences (“edits”) between the read and the reference sequence as shown in the reported alignment. At all distances, HISATx2, STARx2, HISAT align the greatest number of reads, followed by TopHat2.

**Figure 6.**
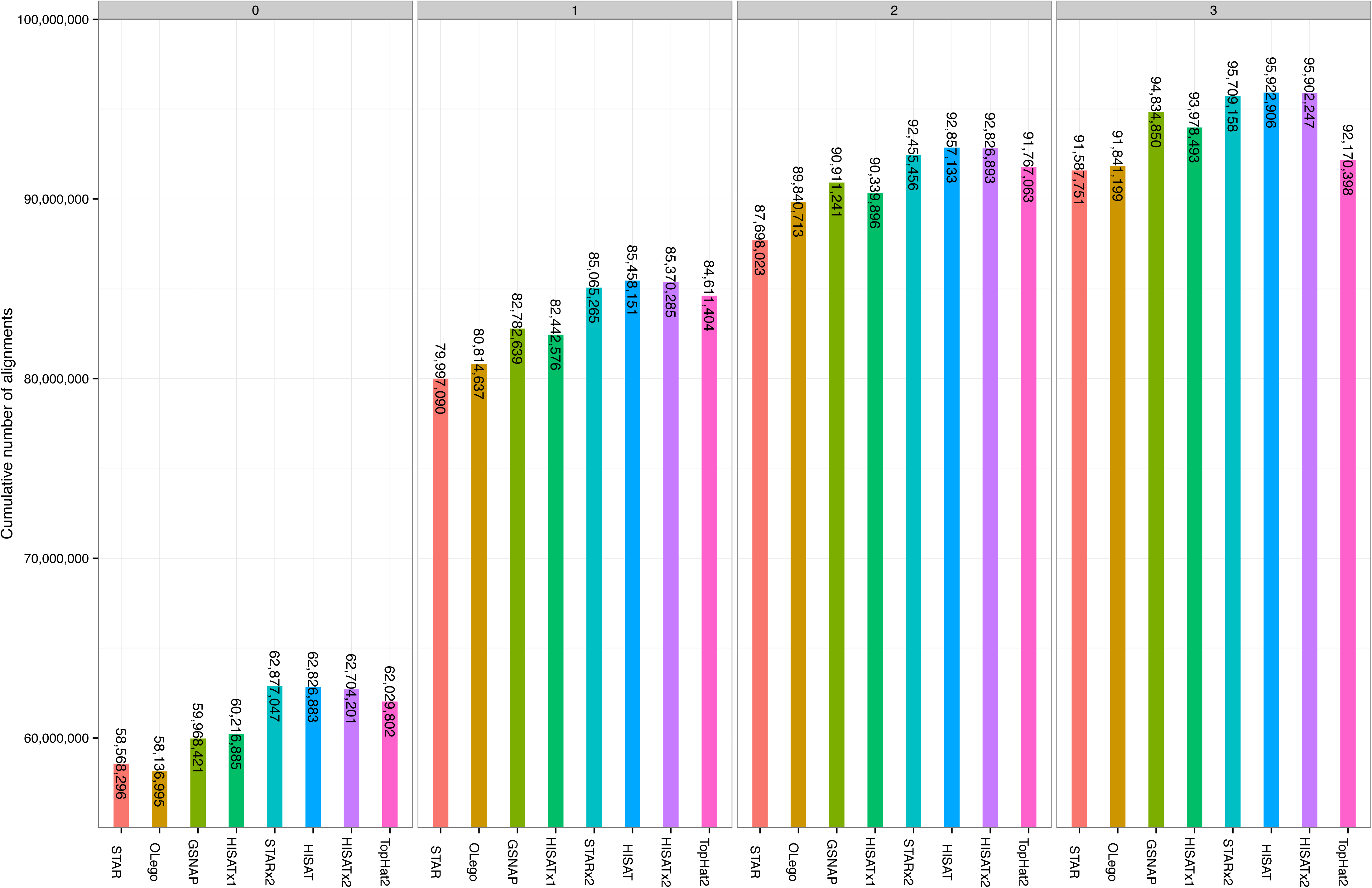
Alignment results for 109 million reads, each 101-bp long, from a human sample. Shown are the cumulative numbers of alignments up to a given edit distance. The leftmost panel shows reads that matched exactly (with an edit distance of 0). The next panel (labelled “1”) shows the number of reads that aligned with either 0 or 1 mismatches; similarly for the panels labelled 2 and 3. Note that GSNAP and STAR report soft-clipped alignments where bases on the ends of reads are left unaligned. To compute edit distances for these alignments, we re-aligned the soft-clipped bases to their corresponding locations in the reference genome and calculated the number of mismatches.

Figure 7 shows the cumulative number of spliced alignments from this same data set that correspond to annotated human splice sites. These are also separated according to edit distance. At every distance and for the overall total, HISATx2, STARx2, and HISAT found the highest numbers of alignments, ranging from 34.6 to 35.1 million. STAR and OLego found the lowest numbers of spliced alignments, at just 26.9 and 26.2 million respectively.

**Figure 7.**
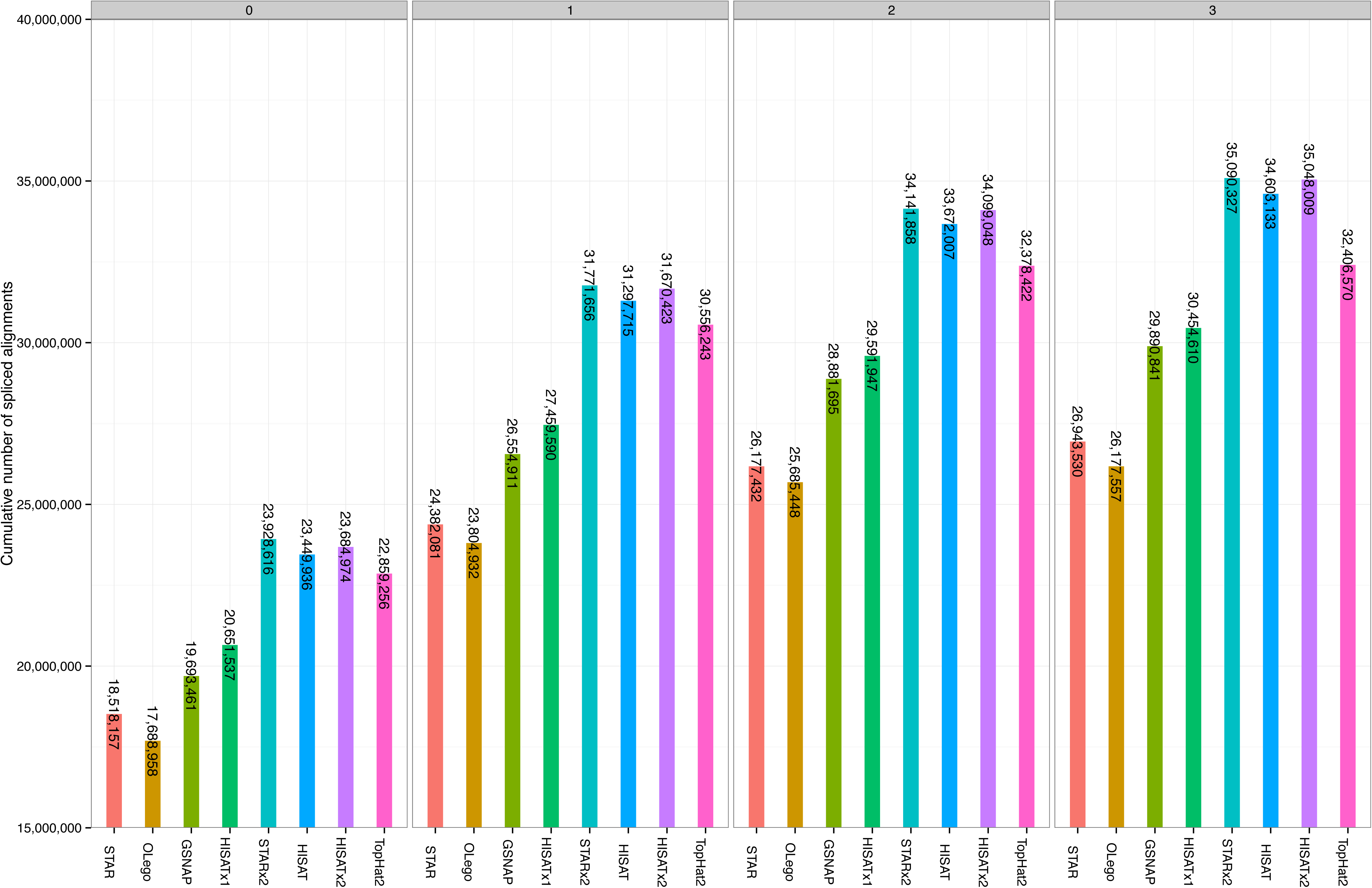
Alignment results of spliced alignment software for 109 million real reads (101-bp long). This figure shows the cumulative number of spliced alignments up to a given edit distance (0, 1, 2, and 3) whose splice sites are known in gene annotations.

Table 2 shows the run-times and memory usages for each program on the 109 reads from the lung fibroblast data set. HISATx1 and HISAT took 23 and 27 minutes (resp.) to process all reads, and only STAR ran in a comparable amount of time, at 24.5 minutes. In contrast, TopHat2 took 1,170 minutes, OLego 990 minutes, and GSNAP 292 minutes. In terms of memory usage, the suffix-array methods STAR and GSNAP used 28 and 20.2 GB of RAM. The BWT-based programs (HISATx1, HISAT, HISATx2, OLego, and TopHat2) required far less memory, ranging from 3.7 to 4.3 GB of RAM.

**Table 2.**
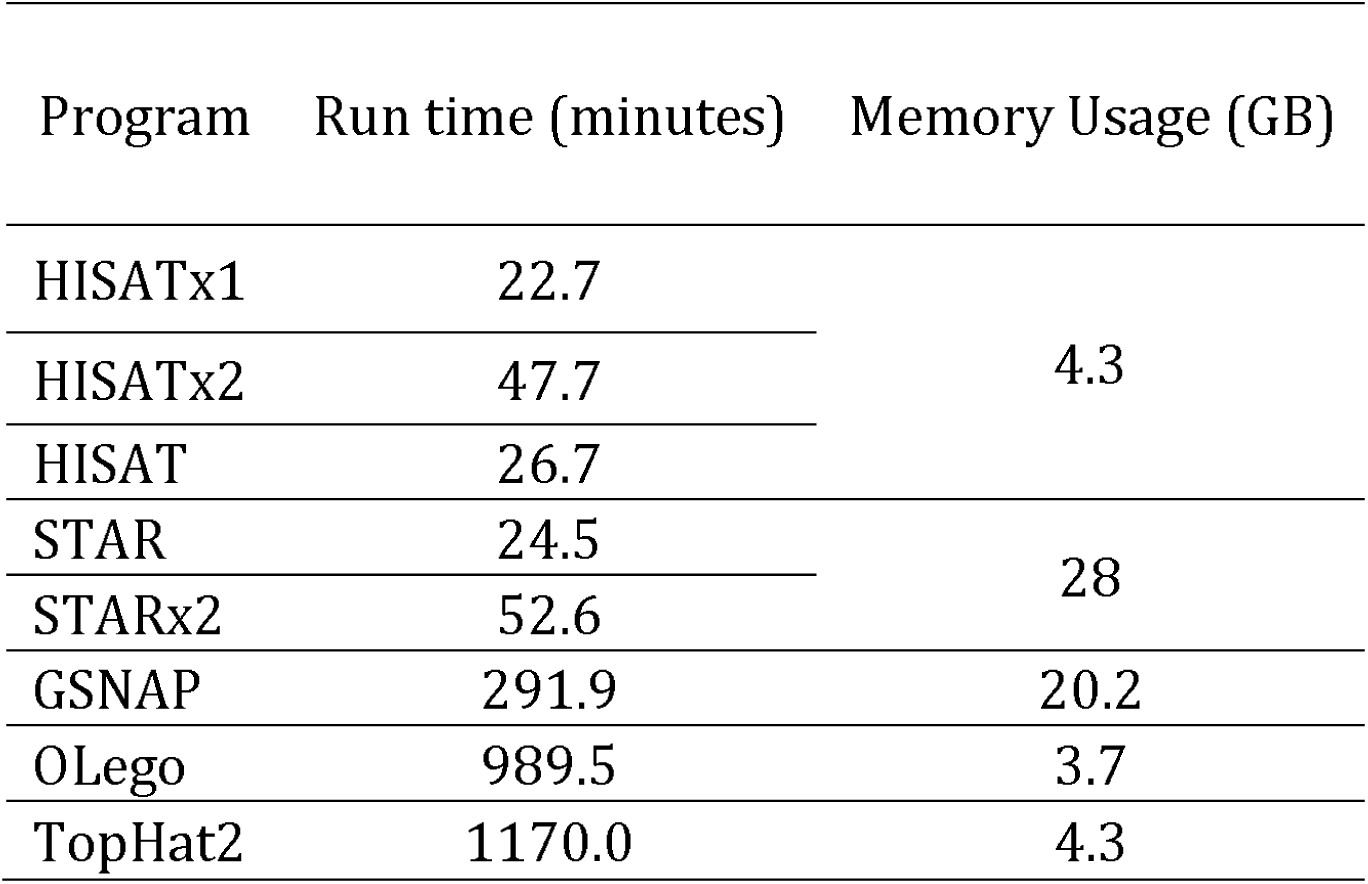
Run times and memory usage for HISAT and other spliced aligners to align 109 million 101-bp RNA-seq reads from a lung fibroblast data set (see main text). We used three CPU cores to run the programs on a Mac Pro with a 3.7 GHz Quad-Core Intel Xeon E5 processor and 64 GB RAM.

Overall, in terms of both alignment quality and speed, HISAT demonstrated superior performance compared to the other programs we tested. In the Supplementary material (Figures S1-S6 and Tables S2-S5), we provide alignment results for additional sets of simulated reads and for an additional real data set from Chen et al.^17^, containing 217 million paired-end reads. In all cases, the relative performances of the alignment programs remained the same as described above. Table S6 also provides details of the input parameters and version numbers for all programs used in these evaluations.

## Conclusions

One of the main purposes of using RNA-seq data is to identify which genes and which isoforms of those genes are expressed in a given sample. Most such analyses begin with an alignment step that maps reads against a reference genome. After mapping the reads, one can group them into loci and then attempt to assemble the reads to reconstruct full-length transcripts and quantify their expression levels. The mapping step is a crucial prerequisite for these analyses, and alignments that are missed cannot usually be recovered at later steps. Because of the large size of RNA-seq data sets today, the alignment of these reads is very time-consuming, taking days or even weeks of compute time.

Most spliced alignment programs use a single global index (e.g., a Bowtie2 index of the human genome) as the basis for alignment. As we have shown, the use of a global index can be very time-consuming when mapping reads with short- and intermediate-length anchors (≤ 15 bp) on one of the exons to which they align. For instance, GSNAP and OLego devote a disproportionate amount of computational time to aligning such reads, while STAR spends less time but finds alignments for only a fraction of them. Also worth noting is that STAR requires ~28 GB of memory, the most of any of the programs tested, due to its use of a suffix array index of the entire genome.

In order to create an aligner that is very fast, very sensitive, and that uses a modest amount of memory, we have designed our new hierarchical indexing scheme and implemented it in HISAT, which achieves these goals by making use of tens of thousands of local indexes, and that gains additional sensitivity from alignment strategies specifically designed to handle different types of reads. These additional indexes combined with the global index enable dramatically faster alignment while matching or exceeding the sensitivity achieved by the best previous spliced aligners. On our real data sets, HISAT was approximately 50 times faster than TopHat2, 42 times faster than OLego, and 12 times faster than GSNAP, with better alignment quality as well. Our experiments on simulated data demonstrate that HISAT has better alignment sensitivity than TopHat2 or any other program. HISAT also has much lower memory requirements (4.3 GB for the human genome) than STAR (28 GB) or GSNAP (18 GB), meaning that HISAT can be run on conventional desktop computers.

In conclusion, HISAT provides better alignment accuracy than TopHat2 and GSNAP, two of the most widely-used spliced alignment programs. It is many times faster than most programs and is slightly faster than STAR, which until now has been the fastest program for this task. While STARx2 and HISAT have similar accuracy, HISAT achieves its performance much more quickly while using much less memory, through the use of local indexes, which are inherently well-suited for aligning across introns. With its combination of high accuracy, speed, and low memory usage, HISAT can produce faster and more accurate results on the very large-scale data sets that are generated in current RNA-seq experiments.

## Materials and methods

HISAT uses its hierarchical indexing algorithm along with several adaptive strategies, based on the position of a read with respect to splice sites, which we describe below. To begin processing each read, it first tries to find candidate locations across the target genome from which the read may have originated. It identifies these locations by first mapping part of each read using the global FM index, which in most cases identifies one or a small number of candidates (Figures 8 and S6). HISAT then selects one of the ~48,000 local indexes for each candidate and uses it to align the remainder of the read. For reads sequenced in pairs, each mate is separately aligned and the alignments of both mates are combined. If a read fails to align, then the alignments of its mate are used as anchors to map the unaligned mate. The extension of each alignment uses an efficient local-index based search as explained below.

**Figure 8.**
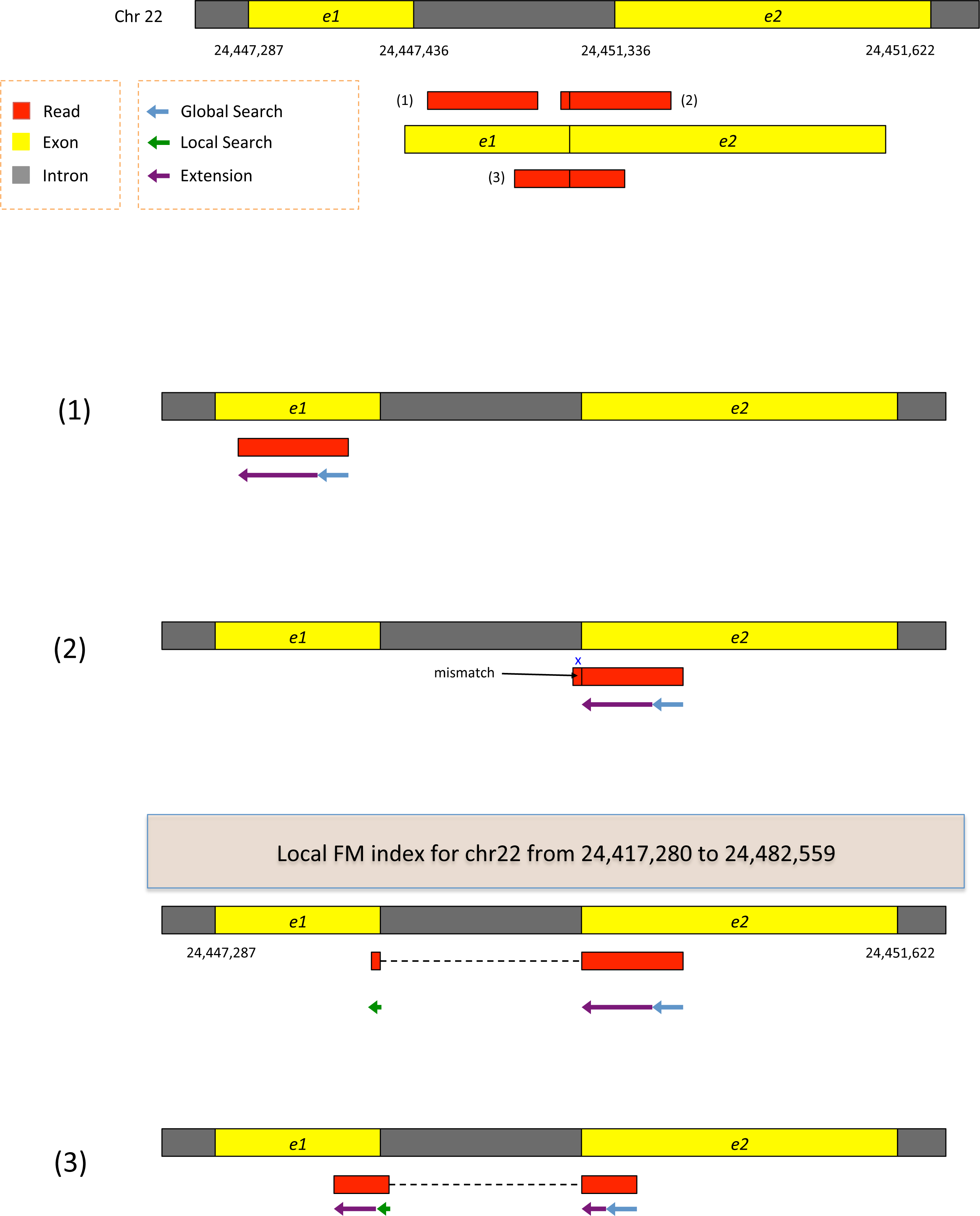
Three working examples demonstrating how HISAT applies its hierarchical indexing for fast and sensitive alignment. The examples include alignment of one exonic read and two junction reads (one an intermediate-anchored read and the other a long-anchored read). Reads are error-free and 100-bp long. See main text for details.

Although searching the global FM index is much faster in principal than k-mer based (or hashing) search, in practice it tends to be slower due to properties of the low-level memory management strategy in a modern computer. The core memory includes both random access memory (RAM) and cache memory, with cache being much smaller but also much faster. When retrieving a block of data, the operating system searches cache first, and only looks in RAM if the block is not found in cache.

Search through a global FM index of the human genome suffers from many cache “misses” because the alignment algorithm proceeds one base at a time through a read and the corresponding locations in a very large FM index (about 750 MB for the human genome). As a match is extended base by base, the search jumps to completely different regions of the index; i.e., the FM index is not organized according to the sequence of the genome itself. Each time the search jumps, the computer has to search RAM and bring in a new piece of the FM index, which is rarely present in the cache already.

In contrast, the far smaller size of our local indexes, 42 KB, allows the entire index to fit in cache, which means that search through the local indexes generates considerably fewer cache misses and therefore runs much faster.

In addition to its two basic operations (global and local searches), HISAT also uses an even faster operation for alignment extension. This operation, which performs direct comparisons of read sequences with genomic sequences, is used only when we know the genomic location to which a read is being mapped. *The extension operation requires the entire genomic sequence to be loaded into memory for fast access; in the case of the human genome this requires 682 MB*. Strategically combining these three operations can dramatically reduce the use of relatively slow operations such as global search and even local search.

Here, we present three different alignment strategies based on three groupings of reads:

1. reads that map either within an exon (M), across 2 exons with at least 15 bp in each exon (2M_gt_15), or across 2 exons with 8-15 bp mapping to one exon (2M_8_15);
2. reads that map to two exons with just 1-7 bp in one exon (2M_1_7) or that map across more than 2 exons (gt_2M); and
3. reads that are likely to be incorrectly mapped to processed pseudogenes.

We illustrate each alignment strategy using examples (error-free reads) that for the purposes of illustration are relatively simple, but that still provide insight into how hierarchical indexing enables fast and sensitive alignment. Although we use error-free reads in these examples, HISAT easily handles reads with both mismatches and indels (see the Supplementary Material and Figure S7 for details). Note that HISAT is optimized for reads ranging from 75-150 bp, the most commonly used (and least expensive) type, but it will also handle the 250-300 bp reads generated by MiSeq instruments.

All the strategies that use local indexes initially just retrieve one index, based on the location of the current match. Among the 246,208 introns from the annotated proteincoding genes in the human genome, 222,503 (90.4%) are completely included in one local HISAT index, each of which spans 64,000 bp. One local index, therefore, is almost always sufficient to align a read. When reads involve long introns, HISAT uses two or more local indexes, up to a maximum intron length of 500,000 bp.

For the examples here, we search for matches in one direction, from right to left, in order to minimize HISAT’s memory footprint, currently 4.3 GB. (Bidirectional search using our method would require 7.5 GB for the human genome.) This unidirectional search does not affect alignment sensitivity, (though it might slightly reduce speed).

*Case 1. Alignment of reads that map either within an exon (M), across 2 exons with at least 15 bp in each exon (2M_gt_15), or across 2 exons with 8-15 bp mapping to one exon (2M_8_15)*.

Figure 8 displays two exons from a gene on human chromosome 22, separated by a 3899-bp intron. Suppose the genomic region is transcribed and spliced, and we have three reads sequenced from the resulting transcript: (1) an exonic read, (2) a read spanning two exons with an 8-bp anchor in one exon, and (3) a read spanning two exons with 50 bp in each exon. All the reads are assumed to be error-free and 100-bp long. We can apply hierarchical indexing to align each of these reads rapidly and correctly. We align the first read using the global FM-index (Figure 8, example 1). Because global search is relatively time consuming, we change strategies when the partial alignment meets two conditions: (1) it is at least 28 bp long and (2) it maps onto exactly one location. For the read shown in the figure, the 28 bp sequence on its right end maps uniquely, allowing us to stop the global search operation at that point. From there, we extend the partial alignment by directly comparing the remaining sequence and the corresponding genomic sequence, which we can extract directly from the genome using the mapped location as an index. Because the read is error-free and contained within one exon, the extension operation sweeps across the remaining 72 bp, completing the alignment for the read. Note that we could perform this alignment using the global FM-index, as TopHat2 does, but the combination of global search and local extension is faster.

For the second read, which has a very short 8-bp anchor on the left side, we first try to map the read using global search, moving right to left as follows (see Figure 8, example 2). The first 28 bp on the right end of the read maps uniquely, allowing us to anchor the alignment and halt the global search. We then extend the alignment until we encounter a mismatch at the 93^rd^ base. This mismatch occurs when the alignment extension reaches the intron. At this point we pause the search, retrieve the local FM-index that contains this location, and align the remaining 8 bp using this index. Because the index covers only a small region, in this case we find just one match for the 8-bp segment. Finally, we check whether the two partial alignments (8 bp and 92 bp) are compatible with each other (e.g., in the correct orientation), and then combine them to produce a spliced alignment of the original read.

Note that if we searched for an 8-bp sequence in the global index, we would expect to find an average of ~48,000 matching locations in the human genome (and sometimes many more). Instead of examining 48,000 possible locations, we use one of the local FM indexes, which is expected to contain just one copy of a given 8-bp sequence, on average. This two-stage hierarchical indexing allows us to avoid examining tens of thousands of candidate locations for short anchors, which in turn dramatically speeds up the overall alignment algorithm.

The third read has long anchors (50 bp each) in each exon. We first align the read beginning on the right, using global search as we did before. After the first 28 bp is uniquely mapped, we switch to the extension operation, which further aligns 22 bp and stops after a mismatch at the 51^st^ base. We then choose a local FM-index and perform a local search using the first 8 bp of the remaining part of the read. Once this 8 bp is found (Figure 8, bottom), HISAT again uses the extension operation to align the rest of the read. Note that depending on how many locations to which the 8-bp is mapped, HISAT uses more base pairs to reduce the number of potential locations within 5.

As we can see from these examples, we can combine global search, local search, and directed read extension to achieve rapid yet sensitive alignment. Note that when a read has multiple spliced alignments, HISAT prefers to report alignments that use the canonical GT and AG dinucleotides on the ends of the intron. From any remaining alignments after this filter, it reports the one with the shortest intron length. HISAT provides several parameters with which users can customize its alignment strategy, including adjustable penalties for mismatches, indels, and non-canonical splice sites.

*Case 2. Reads that map to two exons with just 1-7 bp in one exon (2M_1_7) or that map across more than 2 exons (gt_2M)*

Exon-spanning reads sometimes have very small anchors (defined here as 1-7 bp) in one of the exons. Correctly aligning these reads is extremely difficult because a 1-7 bp anchor will align to numerous locations, even in a local FM index. Arguably the most effective approach to align such short-anchored reads is to use splice site information to remove the introns computationally prior to alignment. We can identify and collect splice site locations when aligning reads with long anchors, and then re-run HISAT for the short-anchored reads, as illustrated in Figure 9. This two-step approach is very similar to the two-step algorithm in TopHat2^9^.

**Figure 9.**
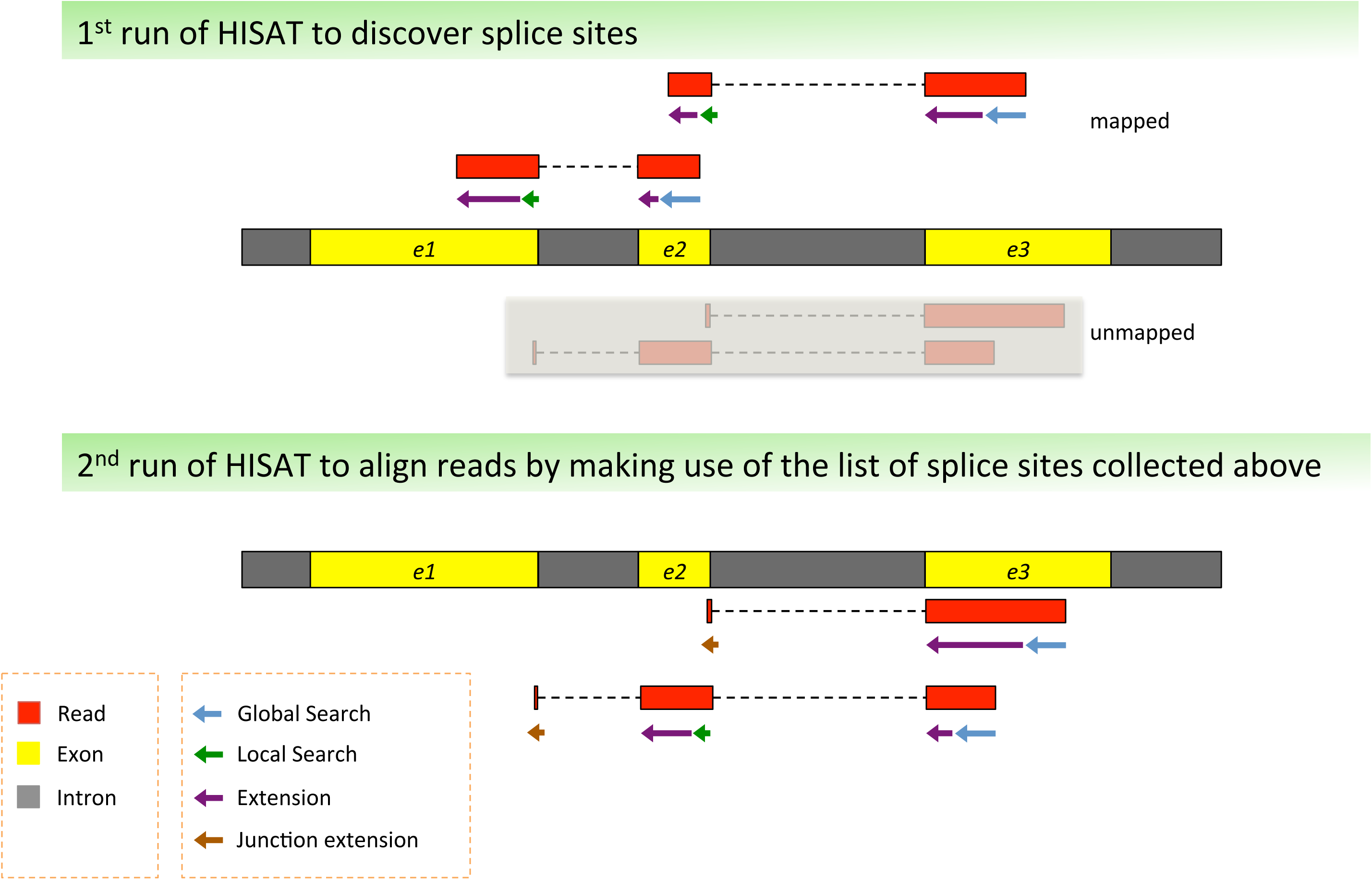
Two-step approach version of HISAT to allow alignment of junction reads with small anchors. This figure shows how to align reads with short anchors (1-7 bp) by making use of splice sites found by reads with long anchors.

More specifically, in the two-step HISATx2 method, we use the first run of HISAT (HISATx1) to generate a list of splice sites supported by reads with long anchors. In the second run we then use the splice sites to align reads with small anchors. For example, consider the unmapped read spanning exons e2 and e3 in the upper portion of Figure 9. The right part of the read will be mapped to exon e3 using the global search and extension operations, leaving a short, 3-bp segment unmapped. We then check the splice sites found in the first run of HISAT to find any splice sites near this partial alignment. In this example, we find a splice site supported by a read spanning exons e2 and e3 with long anchors in each exon. Based on this information, we directly compare the 3-bp of the read and the corresponding genomic sequence in exon e2. If it matches, we combine the 3-bp alignment with the alignment of the rest of the read. This “junction extension” procedure that makes use of previously identified splice sites is represented by brown arrows in Figure 9.

As we show in our experiments on simulated reads (see Results), this two-step strategy produces accurate alignment of reads with very small anchors, as small as 1 bp. Although HISATx2 has considerably better sensitivity, it takes twice as long to run as HISATx1. As an alternative, we developed a hybrid method, HISAT, which has sensitivity almost equal to HISATx2 but with the speed of HISATx1. HISAT collects splice sites as it processes the reads, similarly to the first run of HISATx2. However, as it is processing, it uses the splice sites collected thus far to align short-anchored reads. In the vast majority of cases, it can align even the shortest anchors because it has seen the associated splice sites earlier. This result follows from the observation that most splice sites can be discovered within the first few million reads, and most RNA-seq data sets contain tens of millions of reads. As our results show, HISAT provides alignment sensitivity that very nearly matches the two-step HISATx2 algorithm, with a run-time nearly as fast as the one-step HISAT method.

The hybrid approach is also effective in aligning reads spanning more than two exons, which are more likely to have small anchors. Figure S1 shows that the alignment sensitivity for such reads (gt_2M) increases from 53% using HISATx1 to 95% using HISAT.

*Case 3. Reads that are likely to be incorrectly mapped to processed pseudogenes*

Mis-alignments caused by pseudogenes present additional problems for spliced alignment. Processed pseudogenes are non-functional copies of genes that result when the original gene was transcribed, spliced to remove introns, and re-inserted at a different location in the genome. The most recently created pseudogenes are almost identical to the original genes, meaning that reads from these genes can map equally well to either version of the gene. Intron-spanning reads are a particular problem, because they map end-to-end to the pseudogene, but require a split (spliced) alignment to match the original, active gene. As we showed previously^9^, 2.7% of annotated human genes have pseudogene copies, and the corresponding genes can account for as much as 22.5% of an RNA-seq data set. Therefore pseudogenes can introduce a significant mapping bias unless they are properly handled. Figure 10 illustrates a gene and its corresponding processed pseudogene, where the two exons shown on chromosome 1 have their nearly identical copies on chromosome 17 with only a single base difference. Unlike the two exons on chromosome 1 that are separated by an intron, the two exons on chromosome 17 are adjacent. As a result junction reads originally spanning the two exons on chromosome 1 are likely to be mis-mapped to chromosome 17, particularly if the alignment program prefers contiguous alignments.

**Figure 10.**
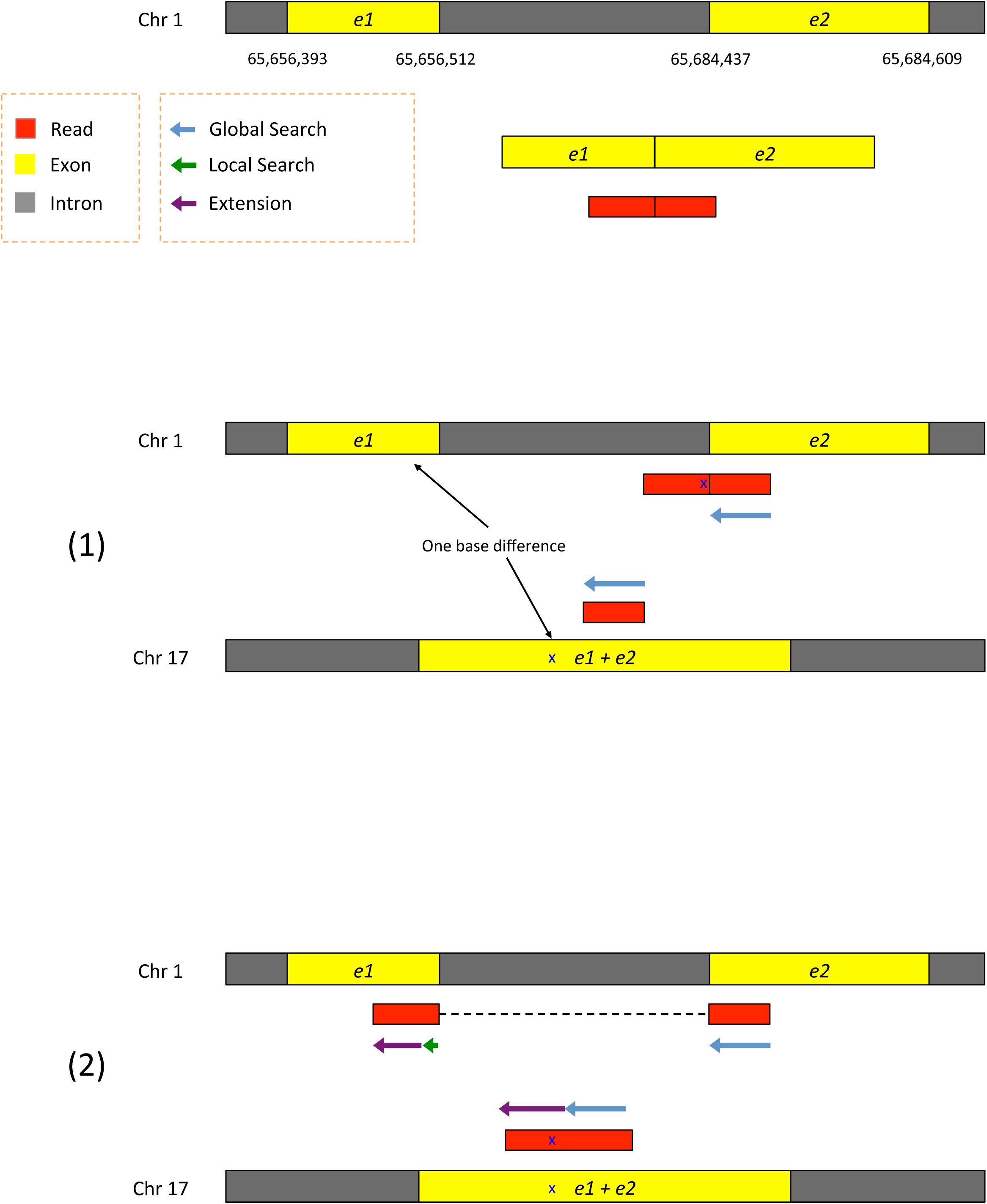
Alignment of junction reads in the presence of processed pseudogenes. This figure shows how to correctly align reads that would otherwise be mapped incorrectly to processed pseudogenes.

As illustrated in Figure 10, HISAT correctly maps reads to their origin by considering several genomic locations. In this example, the rightmost portion of the read (48 bp) maps to chromosomes 1 and 17. The match continues on chromosome 17 (the pseudogene) but is interrupted on chromosome 1 at the 3’ end of the intron. Despite the mismatch, HISAT attempts to extend both partial alignments because both are sufficiently long (at least 22 bp with multiple mappings by default). For the partial alignment on chromosome 1, we resume the search using a local FM index, which yields a spliced alignment with no mismatches. On chromosome 17, the extension of the alignment yields a non-gapped alignment with one mismatch. Given the two candidate alignments, HISAT reports the spliced alignment, because it has no mismatches while the non-spliced alignment has one mismatch. If the two alignments were both equally good, then HISAT would report both alignments. As shown in our results, this alignment strategy allows HISAT to detect more spliced alignments than any of the leading aligners.

## Authors’ contributions

DK, BL, and SLS performed the analysis and discussed the results of HISAT. DK implemented HISAT. DK, BL, and SLS wrote the manuscript. All authors read and approved the final manuscript.

## Acknowledgements

We would like to thank Geo Pertea and Li Song for their invaluable contributions to our discussions on HISAT. We would also like to thank Cole Trapnell for allowing us to use his simulation program (TuxSim). This work is supported in part by the National Human Genome Research Institute (NIH) under grants R01-HG006102 and R01-HG006677 to SLS.

## Competing financial interests

The authors declare no competing financial interests.

## References

1. Mortazavi, A., Williams, B.A., McCue, K., Schaeffer, L. & Wold, B. Mapping and quantifying mammalian transcriptomes by RNA-Seq. Nat Methods 5, 621-628 (2008).

2. Trapnell, C. et al. Differential gene and transcript expression analysis of RNA-seq experiments with TopHat and Cufflinks. Nat Protoc 7, 562-578 (2012).

3. Affymetrix, E.T.P. & Cold Spring Harbor Laboratory, E.T.P. Posttranscriptional processing generates a diversity of 5′-modified long and short RNAs. Nature 457,1028-1032 (2009).

4. Cabili, M.N. et al. Integrative annotation of human large intergenic noncoding RNAs reveals global properties and specific subclasses. Genes Dev 25, 1915-1927 (2011).

5. Kim, D. & Salzberg, S.L. TopHat-Fusion: an algorithm for discovery of novel fusion transcripts. Genome Biol 12, R72 (2011).

6. Garber, M., Grabherr, M.G., Guttman, M. & Trapnell, C. Computational methods for transcriptome annotation and quantification using RNA-seq. Nat Methods 8, 469-477(2011).

7. Grant, G.R. et al. Comparative analysis of RNA-Seq alignment algorithms and the RNA-Seq unified mapper (RUM). Bioinformatics 27, 2518-2528 (2011).

8. Engstrom, P.G. et al. Systematic evaluation of spliced alignment programs for RNA-seq data. Nat Methods 10,1185-1191 (2013).

9. Kim, D. et al. TopHat2: accurate alignment of transcriptomes in the presence of insertions, deletions and gene fusions. Genome Biol 14, R36 (2013).

10. Wu, T.D. & Nacu, S. Fast and SNP-tolerant detection of complex variants and splicing in short reads. Bioinformatics 26, 873-881 (2010).

11. Dobin, A. et al. STAR: ultrafast universal RNA-seq aligner. Bioinformatics 29, 15-21 (2013).

12. Burrows, M. & Wheeler, D.J. A Block-sorting Lossless Data Compression Algorithm. Technical Report 124. Palo Alto, CA: Digital Equipment Corporation (1994).

13. Ferragina, P. & Manzini, G. Opportunistic data structures with applications. Foundations of Computer Science, 2000. Proceedings. 41st Annual Symposium on (2000).

14. Langmead, B. & Salzberg, S.L. Fast gapped-read alignment with Bowtie 2. Nat Methods 9, 357-359 (2012).

15. Wu, J., Anczukow, O., Krainer, A.R., Zhang, M.Q. & Zhang, C. OLego: fast and sensitive mapping of spliced mRNA-Seq reads using small seeds. Nucleic Acids Res 41, 5149-5163 (2013).

16. Griebel, T. et al. Modelling and simulating generic RNA-Seq experiments with the flux simulator. Nucleic Acids Res (2012).

17. Chen, R. et al. Personal omics profiling reveals dynamic molecular and medical phenotypes. Cell 148,1293-1307 (2012).

